# Royal Jelly Derived Extracellular Vesicles Modulate Microglial Nanomechanics and Inflammatory Responses

**DOI:** 10.1101/2025.03.03.641064

**Authors:** Gabriela Zavala, Pablo Berríos, Felipe Sandoval, Graciela Bravo, Nelson P. Barrera, Jessica Alarcón-Moyano, Paulo Díaz-Calderón, Sebastian Aguayo, Christina M.A.P. Schuh

## Abstract

**BACKGROUND:** Microglia, the braińs resident immune cells, undergo profound mechanical and functional changes upon activation contributing to neuroinflammation, a pathological signature of many neurological diseases. Thus, new anti-inflammatory treatment options are needed that tackle these mechanobiological alterations in microglia, which remain strongly understudied. In this context, extracellular vesicles (EVs) are crucial mediators of intercellular and interkingdom communication, yet their influence on the mechanobiological properties of recipient cells remains largely unknown. Honeybee-derived Royal Jelly EVs (RJEVs) have demonstrated remarkable anti-inflammatory properties, but their impact on microglial cellular nanomechanics and uptake mechanisms remains unclear.

**RESULTS:** In this study, we used a multi-disciplinary approach to analyze the resulting biological and nanomechanical changes following the activation of human microglia and the potential effect of RJEV treatment on these mechanobiological parameters. We observed that LPS treatment was associated with decreased cellular Young’s modulus, increased membrane fluidity, and enhanced motility of microglia, indicating a more migratory and pro-inflammatory phenotype. Additionally, LPS exposure altered cellular EV uptake mechanisms by shifting preference from an equilibrium of four mechanisms to the predominance of macropinocytosis and clathrin-dependent endocytosis. Remarkably, RJEV treatment counteracted these mechanobiological changes by, in turn, increasing microglial stiffness, reducing motility, and decreasing secretion of pro-inflammatory cytokines.

**CONCLUSION:** This is the first study to demonstrate that microglial activation state dictates EV uptake mechanisms and to establish a direct link between inflammation, cellular and membrane mechanics, and EV-mediated modulation. Our findings highlight RJEVs as promising candidates for regulating neuroinflammation by targeting microglial mechanobiology as well as opening new strategies for EV-based therapeutics.

**GRAPHICAL ABSTRACT:** 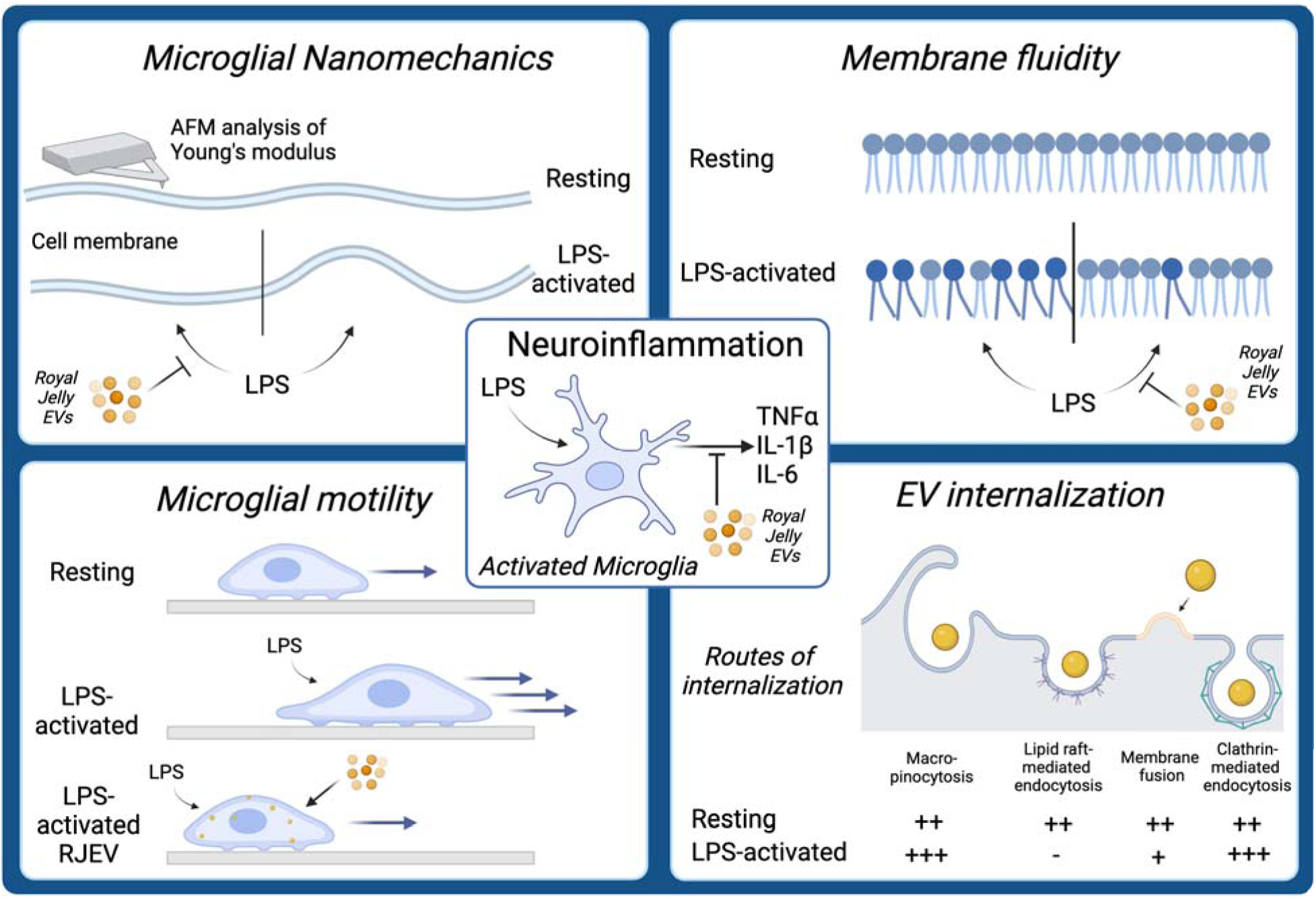

## BACKGROUND

In the last decades, two main discoveries have reshaped our understanding of tissues. The first was the discovery of extracellular vesicles (EVs) as one of the most important factors in cellular communication. EVs are lipid bilayer nanovesicles (30-200 nm) secreted by cells through an elaborate and intricate process, and that contain both active membrane components (such as integrins or tetraspanins (1)) and a diverse cargo comprising proteins (2) and miRNAs (3). Therefore, EVs have several possibilities to exert effects on cells either i) by fusion with the cell membrane, releasing cargo directly into the recipient cell, ii) ligand/receptor interaction and subsequent receptor-based signaling, and iii) internalization into the recipient cell via different types of endocytosis. These routes of entry or cell-interaction are, therefore, defining the subsequent biological effect of EVs on their target cells (summarized by (4)). Most importantly, recent investigations have demonstrated that EV-mediated cellular communication is not limited within one cell type, but can occur between different cell types, species and kingdoms (5).

The second discovery was demonstrating that mechanical forces are as crucial as biochemical signals for tissue development, homeostasis, and cellular functions, which led to the development of the field of mechanobiology (6). The discovery that cells can respond to external mechanical stimuli was followed by findings that cells generate forces through cytoskeletal remodeling and extracellular matrix interactions. Hence, cells are creating feedback loops that influence their mechanical microenvironment, which in turn can promote processes such as cell migration, proliferation, and differentiation in health and disease (summarized by (7)). Even though mechanobiology has gained significant interest in the last decades, so far, the role of EVs in modulating the mechanical properties of cells and tissues has been vastly understudied. One of the cell types that has been reported to undergo significant mechanobiological changes between health and disease is microglia. Upon stimulus (e.g. lipopolysaccharide; LPS), microglia become activated and display changes in migration, actin organization, and traction forces (8,9). Furthermore, they increase proliferation and release oxidants as well as pro-inflammatory cytokines such as Tumor Necrosis Factor α (TNFα), interleukin (IL)1β and IL6, among others (10). This activation of microglia can lead to neuroinflammation, “the activation of the brain’s innate immune system in response to an inflammatory challenge and is characterized by a host of cellular and molecular changes within the brain” (11). Neuroinflammation is a pathological characteristic of a multitude of neurological diseases including classic neuroinflammatory disorders (multiple sclerosis) and neurodegenerative diseases (e.g. Alzheimer’s disease (AD)), but also neuropsychiatric disorders such as schizophrenia (12–15).

Our group recently demonstrated that EVs derived from honeybee Royal Jelly (RJEVs) display anti-inflammatory effects in LPS-stimulated macrophages (16). Moreover, it was found that RJEVs utilize different mechanisms to exert their effects in human cells such as mesenchymal stem cells and fibroblasts, including membrane fusion, and clathrin-mediated endocytosis. (17) Several studies reported anti-neuroinflammatory effects of Royal Jelly (RJ) and its components in both *in vitro* and *in vivo* assays. In an LPS-induced model, RJ inhibited the expression of pro-inflammatory cytokines such as TNF-α, IL-1β and IL6, chemokines (CCL-2, CCL-3), and iNOS (18,19) *in vitro* and *ex vivo*. However, even though anti-neuroinflammatory effects have been observed by several studies, it remains unclear how RJ exerts these effects, especially in light of recent insights into the importance of mechanobiology and EVs.

Therefore, in this study we analyzed in detail how LPS activation changes mechanical properties of human microglia, such as cellular Young’s modulus, motility, and membrane fluidity, and how treatment with RJEVs can revert the observed changes. Overall, this is the first mechanistic study linking cell stiffness, membrane fluidity, and biological responses to changes in the activation state of human microglia.

## METHODS

### Cell culture

Human microglial cells (HMC3) were purchased from ATCC (CRL-3304) and cultivated as indicated by the manufacturer. Cell culture media for cell expansion consisted of Eagle’s Minimal Essential Media (EMEM, Sigma) with 10% Fetal Bovine Serum (FBS, Sigma), 1% Penicillin/Streptomycin (PenStrep, Thermo Scientific) and 1% Sodium pyruvate (expansion media). Cultures were kept at a density of 2-4×10^4^ cells per cm^2^. All experiments were conducted under normoxic conditions and at 37°C (humidified incubator). To induce inflammation, Lipopolysaccharide (LPS) from Escherichia coli O111:B4 was used at a concentration of 1 µg/ml (Sigma).

### Royal Jelly EVs (RJEVs) isolation and characterization

RJEVs were isolated as previously described (16). Royal Jelly was purchased from a local provider in Chile (El Alba). All batches utilized for this study were analyzed with Nanoparticle Tracking analysis to assess size distribution and concentration, as previously described (16).

### Zeta Potential

EV colloidal properties were measured with a ZetaSizer (Malvern Panalytical, Malvern, UK. Pro Red model) using dynamic light scattering analysis (DLS) at room temperature. Samples were loaded into a Folded Capillary Zeta Cell at a 1:10 dilution with ddH_2_O. A refraction index of 1.46 was used for RJEVs in ultrapure water as a dispersant. The samples were measured in technical triplicate and analyzed with the ZS Xplorer software.

### Quantification of Total Phospholipid Content

Phospholipid enrichment in RJEVs was quantified using a “Phospholipid assay kit” (Sigma Aldrich) according to the manufacturer’s instructions and as described in ((20), supplement). Absorbance was measured at 530 nm with a microplate reader (Tecan Sunrise, Austria). RJEV isolates were matched with RJ raw material.

### Quantification of Total Carbohydrate Content

Carbohydrate impurities were assessed with the “Total Carbohydrate Assay Kit” (Sigma Aldrich) according to the manufacturer’s instructions and as described in ((20), supplement). Absorbance was measured at 490 nm using a microplate reader (Tecan Sunrise).

### Microglia cell viability and proliferation

The viability of HMC3 microglial cells was determined with measuring metabolic activity (MTT assay) and presence of lactate dehydrogenase (LDH assay). Cells were seeded in duplicates in a 96-well plate with media containing 5% FBS (1×10^4^ cells per cm2) and left to adhere overnight. The next day media was changed to 2.5% FBS (125 µl). LPS and RJEVs at different concentrations (0.001, 0.01, 0.1, 0.5, 1, 5, 10 µg) were added. Untreated cells and 2.5% Triton-X 100 served as controls. After a 24-hour incubation period, 60 µl supernatant from each well were transferred to a new plate and kept at 4°C for the LDH assay. Cells with the remaining media were incubated with 500 µg/ml 3-(4,5-dimethylthiazol-2-yl)-2,5-diphenyltetrazolium bromide (MTT reagent, Sigma) for 1 hour at 37°C. Afterward, the media was discarded, and precipitated formazan was dissolved in 75 µl dimethyl sulfoxide (DMSO, Sigma). Absorbance was measured at 570 nm with a microplate reader (Tecan Sunrise, Tecan). LDH assay was performed according to manufacturer’s instructions (Cytotoxicity detection Kit (LDH), Roche, 11644793001). Briefly, supernatants were brought to room temperature, and the LDH reaction mixture was added to the supernatants at a ratio of 1:1 for 30 minutes. Reaction was stopped with 1N hydrochloric acid (HCl) and absorbance was measured at 492 nm.

On the other hand, proliferation was measured with a BrdU assay (Roche) according to the manufacturer’s instructions. Cells were seeded as described above. Additional to LPS and RJEVs, the BrdU labeling solution was added. After 24 hours, supernatants were discarded and cells were fixed and denatured with FixDenat solution. Subsequently, wells were incubated with anti-BrdU-POD solution for 90 minutes. Wells were washed with wash solution and reaction was developed with substrate solution for 20 minutes. Reaction was stopped with 1N HCl and absorbance was measured at 492 nm.

### ELISA of pro-inflammatory cytokines

The anti-inflammatory potential of RJEVs on microglia was assessed after stimulation with LPS by analyzing the subsequent secretion of pro-inflammatory cytokines. Cells were seeded at a density of 8×10^3^ cells/cm^2^ in 6-well plates and left to adhere overnight. RJEVs were added in concentrations of 1, 10, 100, 1000 and 2500 EVs per cell 4h, and 20 µg/ml indomethacin (inhibition control, non-steroidal anti-inflammatory drug) 2h prior to stimulation with LPS. After 24h LPS stimulation, the supernatant was collected and centrifuged at 2000xg for 20 min to remove cellular debris. Tumor Necrosis Factor alpha (TNFα), Interleukin 1 beta (IL-1β) and Interleukin 6 (IL-6) were quantified with sandwich ELISA (DuoSet, R&D Systems) according to manufacturer’s instructions. Results were normalized to 100.000 cells.

### F-actin visualization

HMC3 cells were seeded onto 12 mm glass coverslips at a density of 7.5×10^3^ cells/cm^2^. After the respective stimulus, cells were washed and fixed with 4% paraformaldehyde (PFA) at room temperature for 10 minutes. Cellular F-actin was stained with phalloidin (Alexa Fluor 488-conjugated phalloidin, Thermo Fisher, US) at a concentration of 165 nM in staining buffer (PBS containing 1% BSA and 0.2% Triton X-100). After incubation for 1 h at RT protected from light, cells were washed with PBS. Cell nuclei were counterstained with HOECHST 33342 (10 µg/ml, Thermo Fisher) for 5 minutes. Subsequently, coverslips were washed once with PBS, mounted onto glass slides with Fluorsave (Merck Millipore, US) and left to dry at 4°C in the dark overnight. Samples were analyzed with the Fluoview FV10i confocal microscope and processed with the Olympus Fluoview V4.2b software.

### Routes of RJEV internalization into microglia

An uptake inhibition assay was performed in order to identify the routes of RJEV internalization into microglia, by utilizing an adaptation of a previously described for human mesenchymal stem cells (16). RJEVs were stained with 5 (6)-Carboxyfluorescein diacetate N-succinimidyl ester (Cell Trace CFSE, Thermo Fisher Scientific) and subsequently quantified with NTA (21). 5×10^4^ HMC3 microglial cells were seeded into 6-well plates and the next day, cells were incubated with 1 µg/ml LPS. After 24 hours, one well per condition (ctrl, LPS) was counted. Potential routes of uptake were blocked with Amiloride 50 µM, Chlorpromazine 28 µM or Omeprazole 1 mM for 1 hour and Filipin 7.5 µM (for 15 min). Subsequently, 250 CFSE-RJ-EVs were added per cell and incubated for 4 hours. Cells were washed 2x with PBS, detached with accutase and analyzed with flow cytometry (FACS Canto II, Becton Dickinson). The FlowJo software (TreeStar, version 8.8.6) was used for data analysis.

### Cell motility measurement

Motility of HMC3 microglia cells was measured with a Phio CellWatcher M (Phio, Germany). 1×10^4^ cells were seeded in 6 well plates and left to adhere for 24h. The next day, cells were washed twice with 1.5ml PBS and the media was changed to 1% FBS (1ml, exact). After 24 hours with low-FBS media, LPS and RJEVs were added at different concentrations (100, 1000 and 2500/cell). Untreated cells and cells incubated with RJEVs or LPS alone served as controls. The plate was placed into the CellWatcher microscope in a humidified incubator at 37°C and 5% CO_2_. Images were taken every 30 min for 24 hours. Analysis was divided into five different time sections: 0-4.5h (initial phase), 5-9.5h, 10-14.5h and 15-19.5h (mid phases) and 20-24h (late phase).

### Atomic force microscopy (AFM) live-cell nanomechanics

For living cell AFM nanomechanical experiments, HMC3 were seeded in PLL-treated 12 mm glass coverslips at 7500 cells/cm2 cell density and were left to adhere for a minimum of 24 h. RJEVs were added at 1000 EV/cell ratio for 18-20 h and LPS 1 µg/ml was added for 4 h before the AFM analysis. An MFP 3D-SA AFM (Asylum Research, USA) was employed using TR400PB cantilevers with a nominal spring constant of 0.09 N/m (Asylum Research, UK). Individual tip calibration was performed before each experiment with the proprietary Asylum Research Software GetReal™ Automated Probe Calibration method. With the goal of extracting cell elasticity, each calibrated cantilever was carefully approached toward the surface of an attached individualized cell by means of a CCD camera, to generate a 50×50 µm force map with 32×32 force-curves for a total of 1024 force-curves per scan. Force curves were obtained with a constant velocity of 8 µm/s, a 5 nN maximum force, and a maximum pulling length of 8 µm in the z-axis. All force-curve (FC) measurements were performed in buffer (Hanks balanced salt solution, HBSS, Gibco), for no longer than 2 hours to ensure proper cell viability throughout experimentation. From the resulting live cell stiffness values, Young’s modulus was obtained for each force curve using the Johnson, Kendall, and Robert (JKR) model in the Asylum Research proprietary software (22). Finally, all data and images were obtained and analyzed using the Asylum Research proprietary software and elasticity histograms were generated in IGOR 8.04.

### Membrane motility measurements (Fluorescence recovery after photobleaching, FRAP)

HMC3 were seeded in PLL-treated 22 mm glass coverslips at an initial density of 1×10^4^ cells/cm^2^. Cells were left to adhere overnight. Subsequently, cells were incubated with RJEVs (1000 RJEVs/cell) for 20 h and LPS (1 µg/ml, Sigma Aldrich) was added for 4 h before the analysis. The plasma membrane was stained with 5 µg/ml Cell Mask Green (Thermo Fisher) in cell culture media for 4 minutes at room temperature. Subsequently, the media was replaced, and FRAP measurements were acquired using the LSM 880 Zeiss Airyscan confocal microscope (ambient controlled, 37°C, 5% CO_2_) and the software Zen Black 2.3. After selecting the highest fluorescence intensity plane, cells were bleached with a 514 nm argon laser at 100%. 5 photographs were acquired before the bleaching, and the recovery was followed for 40 sec in total. Frames were taken every 50 ms.

### Statistics

Data in this study are shown as mean ± standard deviation, box plot with whiskers to minimum and maximum, or violin plots with median and 25%/75% percentiles. Normality was tested with Shapiro-Wilk test. Student-t test (for two groups) or one-way ANOVA followed by Tukey’s range test (for three or more groups) were used for normally distributed data. Mann-Whitney and Kruskal-Wallis tests were used for non-parametric data. Analysis was performed using GraphPad Prism 9 for Mac OS X, version 9.3.1 (GraphPad Software, Inc.). Significance was considered at p<0.05 (see figure legends for specific values). N values were determined from independent experiments and independent isolations of RJEVs.

## RESULTS

### Royal Jelly EVs

RJEVs isolated for this study displayed an average yield 1.48×10^10^ ± 2.17×10^9^ particles per gram of RJ. NTA revealed an average particle size of 151.5 nm ± 16.73 nm and a median size 131.9 nm ± 22.43 nm. No statistical difference was found between average and median particle size, indicating a mostly homogeneous EV population (**Figure 1A-B**). The Zeta potential was - 9.01 ± 3.9 mV (**Figure 1C**). RJEVs displayed a round, sphere-like shape (**Figure 1D**). Analysis of carbohydrate impurities demonstrated a significant reduction of carbohydrates in isolated EVs in comparison to the prime material (RJ 5.39±0.45 µg/µl, RJEV 7.9±5.4 ng/µl) (**Figure 1E**). Total phospholipid analysis, an indicator for lipid enrichment, showed a significantly higher lipid content in the isolated RJEVs (167.3 ± 14.89 µM) compared to the prime material (64.45 ± 13.35 µM) (**Figure 1F**).

**Figure 1:**
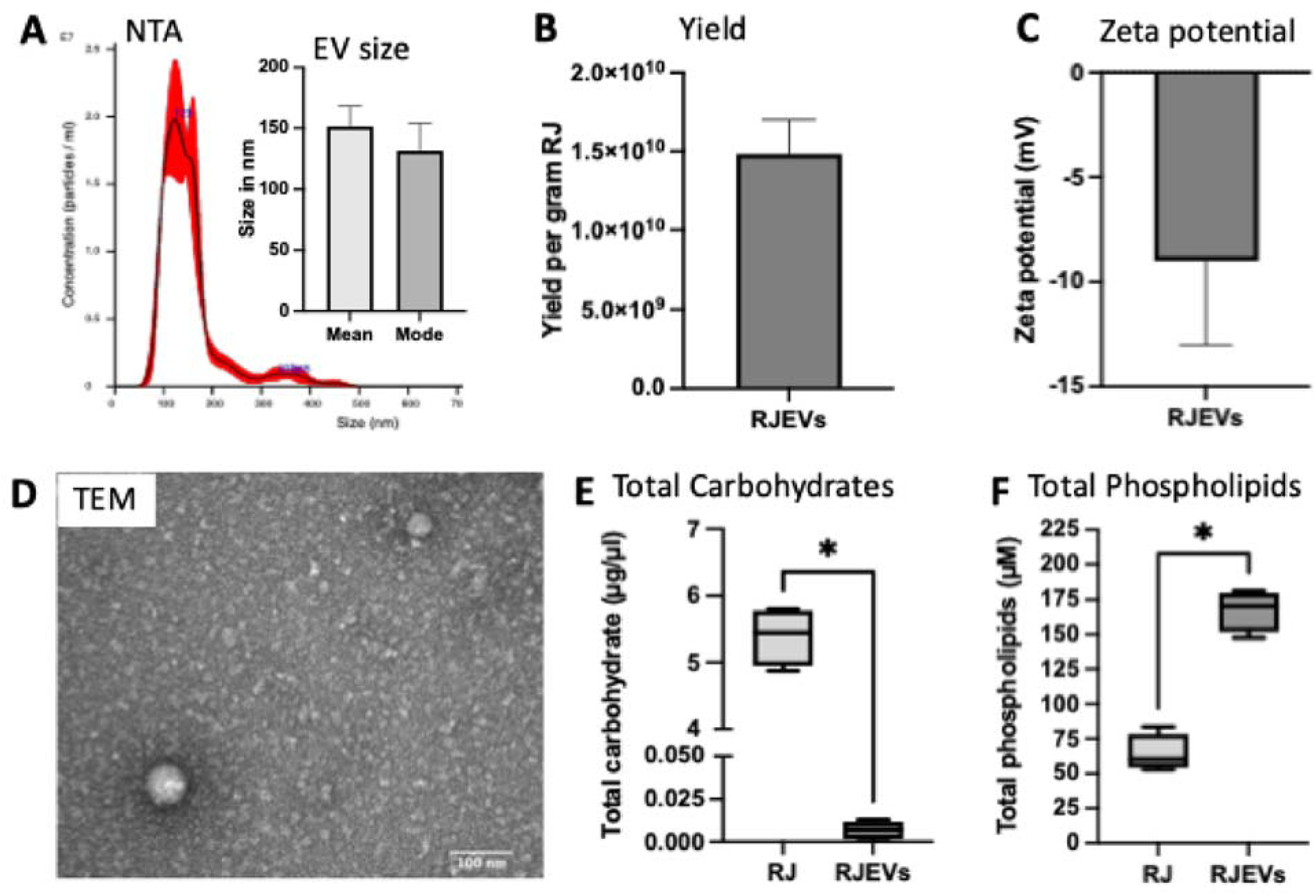
Characterization of RJEVs: A) representative NTA histogram of RJEVs and average (mean) and median (mode) particle size; n=8; B) Particle yield of RJEV per gram raw RJ; n=8; C) Zeta potential of RJEVs in mV, n=5; D) representative TEM micrograph of RJEVs, scale bar shows 100 nm; For A/B/C data is displayed as mean ± SD E) Quantification of total carbohydrates, isolated EVs (RJEVs) were compared to pre-isolation supernatant (RJ); n=4; F) Quantification of total phospholipids, isolated EVs (RJEVs) were compared to pre-isolation supernatant (RJ); n=4; For E/F, data is displayed as box and whiskers (min-max); Statistical significance was tested with Mann-Whitney test; *P<0.05

### Effects of LPS and RJEVs on cellular functions

Metabolic activity, cell damage and proliferation of HMC3 microglia were measured after exposure to LPS and in the presence and absence of different concentrations of RJEVs. LPS significantly increased cell damage (12.4% ± 2.5%; **Figure 2B**) and decreased cell viability (−13.1% ± 3.7%, **Figure 2A**). RJEVs at concentrations between 10 and 2500 EVs/cell increased the metabolic activity of LPS-treated cells to levels above non-LPS-treated controls (4.6% to 8.9%). Cell damage in the presence of 1 to 2500 RJEVs per cell was lower than in the LPS-treated cells (−11.2% to 14.6%). Interestingly, no dose-dependent effect was observed for metabolic activity and cell damage. For cell proliferation (**Figure 2C**), a reverse dose-dependent effect was observed: LPS significantly stimulated proliferation (+31.4% ± 2.8%), which was inhibited by lower dosages of RJEVs (+1.3% to 11.5%), and increased with higher dosages (20.7% to 29.5%). Morphology of the HMC3 microglia can be observed in **Figure 2E**, stained with phalloidin for F-actin.

**Figure 2:**
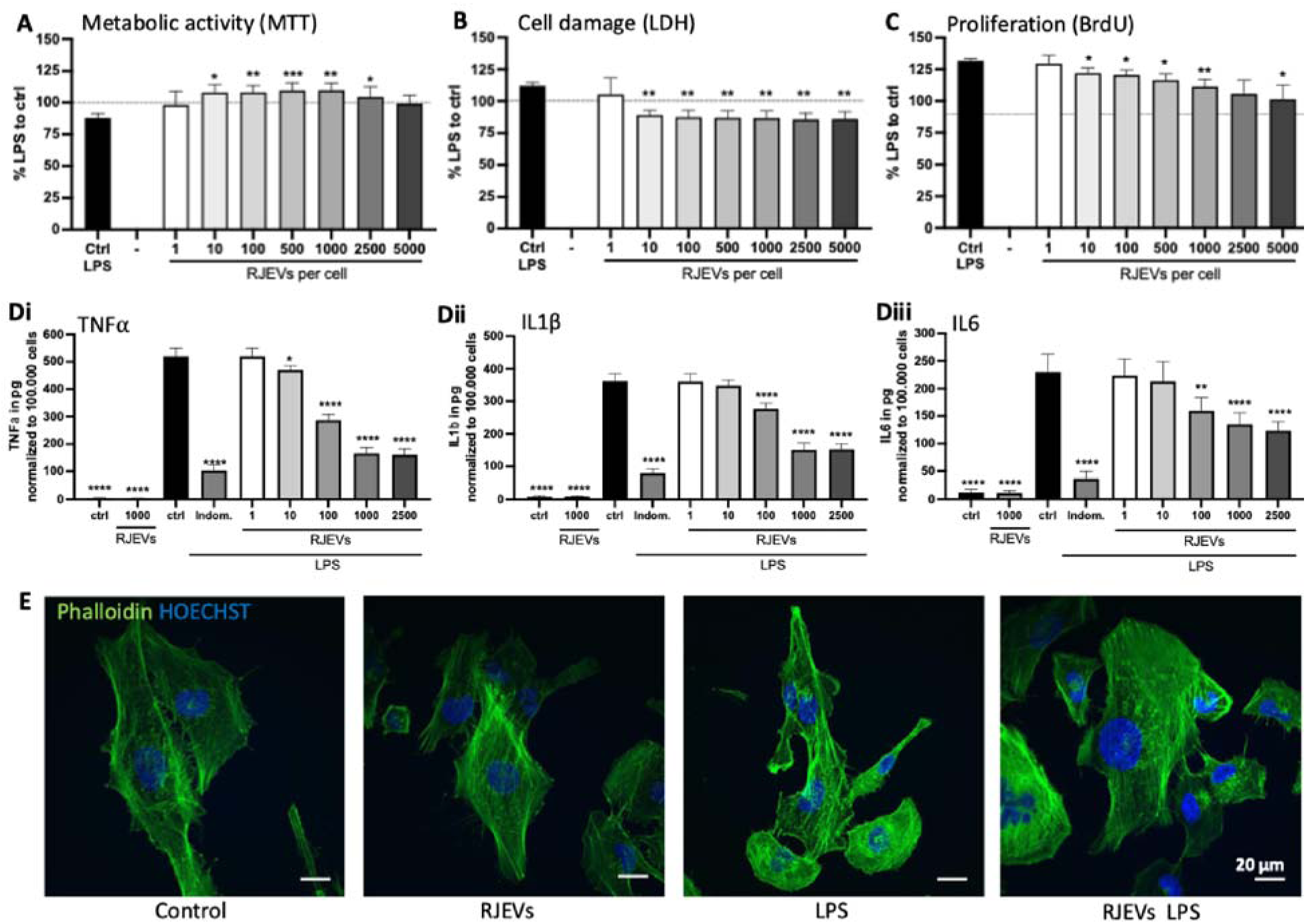
Effect of RJEVs on cellular function and inflammatory responses in HMC3 cells; A) Effect of RJEVs on metabolic activity in LPS-stimulated HMC3 cells, measured with MTT assay; B) Effect of RJEVs on cellular damage in LPS-stimulated HMC3 cells, measured with LDH assay; Effect of RJEVs proliferation in LPS-stimulated HMC3 cells, measured with BrdU assay; n=5; ; For A/B/C data is displayed as percentage of LPS-stimulated cells to control in presence of different concentrations of RJEVs (1-5000 RJEVs per cell); mean ± SD; n=5; D) secretion of pro-inflammatory cytokines IL-1, IL-6, TNFIZI by HMC3 cells after stimulation with LPS, measured with ELISA; controls with and without LPS and in presence of Indomethacin (Indom.); results were normalized on 100.000 cells; n=4; A/B/C/D Significance was tested with one-way ANOVA with Tukey’s post-hoc; *p<0.05, **p<0.01, ***p<0.001, ****p<0.0001; * indicates difference to LPS ctrl; E) representative images of HMC3 cells, stained with Phalloidin (F-actin, green) and HOECHST (nucleus, blue); scale bar represents 20 µm.

### Anti-inflammatory effect of RJEVs in activated microglia

The potential anti-inflammatory effect of RJEVs in LPS-stimulated microglia was assessed with ELISA, quantifying the presence of pro-inflammatory cytokines TNFα, IL1β, and IL6 secreted into cell culture media (**Figure 2D**). Cells not stimulated with LPS expressed neglectable levels of TNFα, IL1β and IL6 (0-15pg). RJEVs did not stimulate an inflammatory response, and cytokine secretion remained at the respective baseline levels. Stimulated with LPS, microglia significantly increased secretion of pro-inflammatory cytokines (TNFα: 519.6 pg ± 30.3 pg; IL1β: 362 pg ± 22.3 pg; IL6: 230.1 pg ± 32.2 pg). The non-steroidal anti-inflammatory drug Indomethacin significantly reduced the secretion of pro-inflammatory cytokines, however not to the baseline level. RJEVs displayed a dose-dependent effect, between 1 and 1000 RJEVs per cell, which did not further increase for 2500 RJEVs per cell. For TNFα, initial effects were observed with 10 RJEVs per cell, and for IL1β as well as IL6 with 100 RJEVs per cell. Means and SD for each condition, as well as significance between each group, can be found in **Supplemental Tables 1 and 2**.

### Microglia change their route of EV internalization after activation

To assess EV internalization, it has been established to utilize fluorescence-labelled EVs. Confocal images in **Figure 3C** demonstrate HMC-3 cells after incubation with CFSE-labelled RJEVs. The four routes of EV internalization (macropinocytosis, membrane fusion, lipid raft-mediated endocytosis, and clathrin-dependent endocytosis) were blocked individually in the presence and absence of LPS. The higher the difference to the control when inhibited, the higher the involvement of the tested mechanism in EV uptake. Treatment with LPS significantly increased the uptake of RJEVs, from 20.28% ± 1.77% to 26.98% ± 1.23% RJEV-positive cells (**Figure 3B**). Then, assessing the mechanisms of internalizations, a major shift was observed when microglia were stimulated with LPS. While the unstimulated control group displayed a balanced involvement of all four mechanisms (macropinocytosis 28.22% ± 4.50%, clathrin-mediated endocytosis 39.88% ± 3.80%, lipid raft-mediated endocytosis 22.04% ± 4.14% and membrane fusion 43.80% ± 4.97% inhibited with the respective inhibitors), LPS significantly shifted the mechanisms toward macropinocytosis and clathrin-dependent endocytosis (53.09% ± 3.75% and 56.49% ± 3.70%, respectively). Membrane fusion still displayed an involvement of around 32.10%; however, lipid raft-mediated uptake inhibition was decreased to 4.21% following LPS exposure (**Figure 3A**).

**Figure 3:**
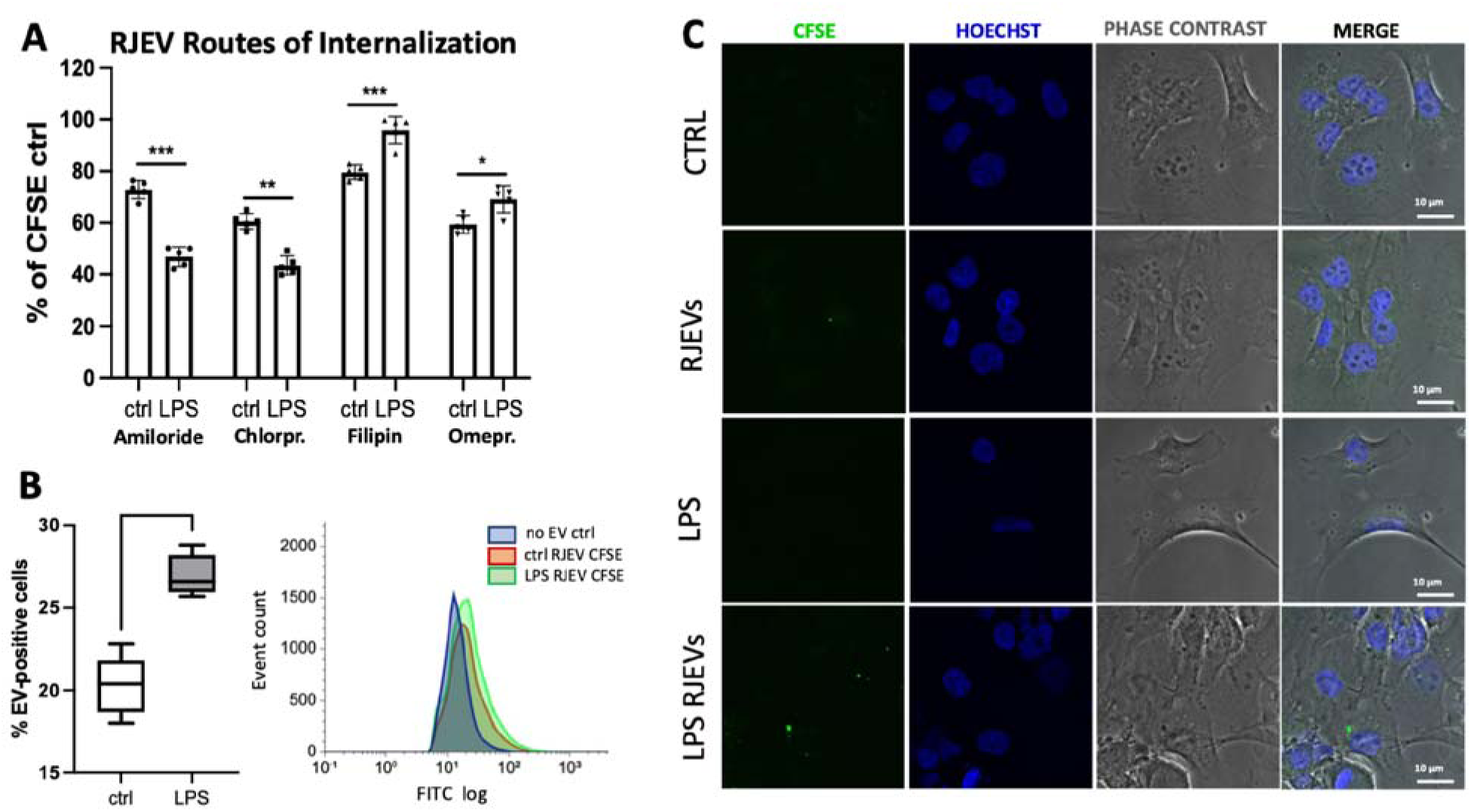
Route of internalization into microglia; A) Quantitative analysis of CFSE-stained RJEV internalization inhibition measured with flow cytometry; Routes of internalization were blocked with amiloride (micropinocytosis), chlorpromazine (clathrin-dependent endocytosis), filipin III (lipid raft-mediated endocytosis) and omeprazole (membrane fusion), in non-stimulated (ctrl) and LPS-stimulated HMC3 cells. N=5; Significance was tested with one-way ANOVA with Tukey’s post-hoc; *p<0.05, **p<0.01, ***p<0.001, ****p<0.0001; Bi) Quantification of RJEV-positive HMC3 cells, in absence (ctrl) and presence of LPS (LPS); n=5; significance tested with student-t test; *p<0.05, **p<0.01; Bii) representative flow cytometry histogram showing HMC3 cells without RJEVs (blue), and with RJEVs in absence (red) and presence of LPS (green). C) Representative confocal images of HMC3 cells in absence and presence of LPS and CFSE-stained RJEVs;

### RJEVs inhibit LPS-induced motility

Subsequently, cell motility was assessed with the Phio CellWatcher by tracking cells over 24h after stimulation with LPS and RJEVs at different concentrations (**Figure 4A and 4B).** HMC3 microglia displayed an overall migration speed of 1.71 µm/h to 2.41 µm/h. RJEVs without LPS stimulation did not show significant differences compared to the untreated control, independent of the time frame. LPS, however, significantly increased motility after 5 hours and continuously until 24 hours. The highest effect of LPS was observed between 5 and 15 hours, with motility slowly decreasing towards 24 hours. Interestingly, RJEVs at concentrations 1000/cell and 2500/cell displayed significantly lower motility from the initial phase on and throughout the experiment. The differences ranged from a 10.7% to 16.4% reduction for 1000 RJEVs/cell and a 19.5% to 24% reduction for 2500 RJEVs/cell. The motility reduction of 100 RJEVs/cell was only significant within the time frame of 10-14.5h, and ranged from 4.8% to 10.7%. Overall, it was observed that motility slightly decreased over time, from 2.8 µm/h in the first time frame (0-4.5h) to 2.4 µm/h in the last time frame (20-24h). The observed growth rate was between 0.4% and 2.4%. The LPS-stimulated group displayed a trend toward higher proliferation however, the changes were not significant (**Figure 4C**). Descriptive statistics and statistical analysis can be found in **Supplemental Tables 3 and 4**.

**Figure 4:**
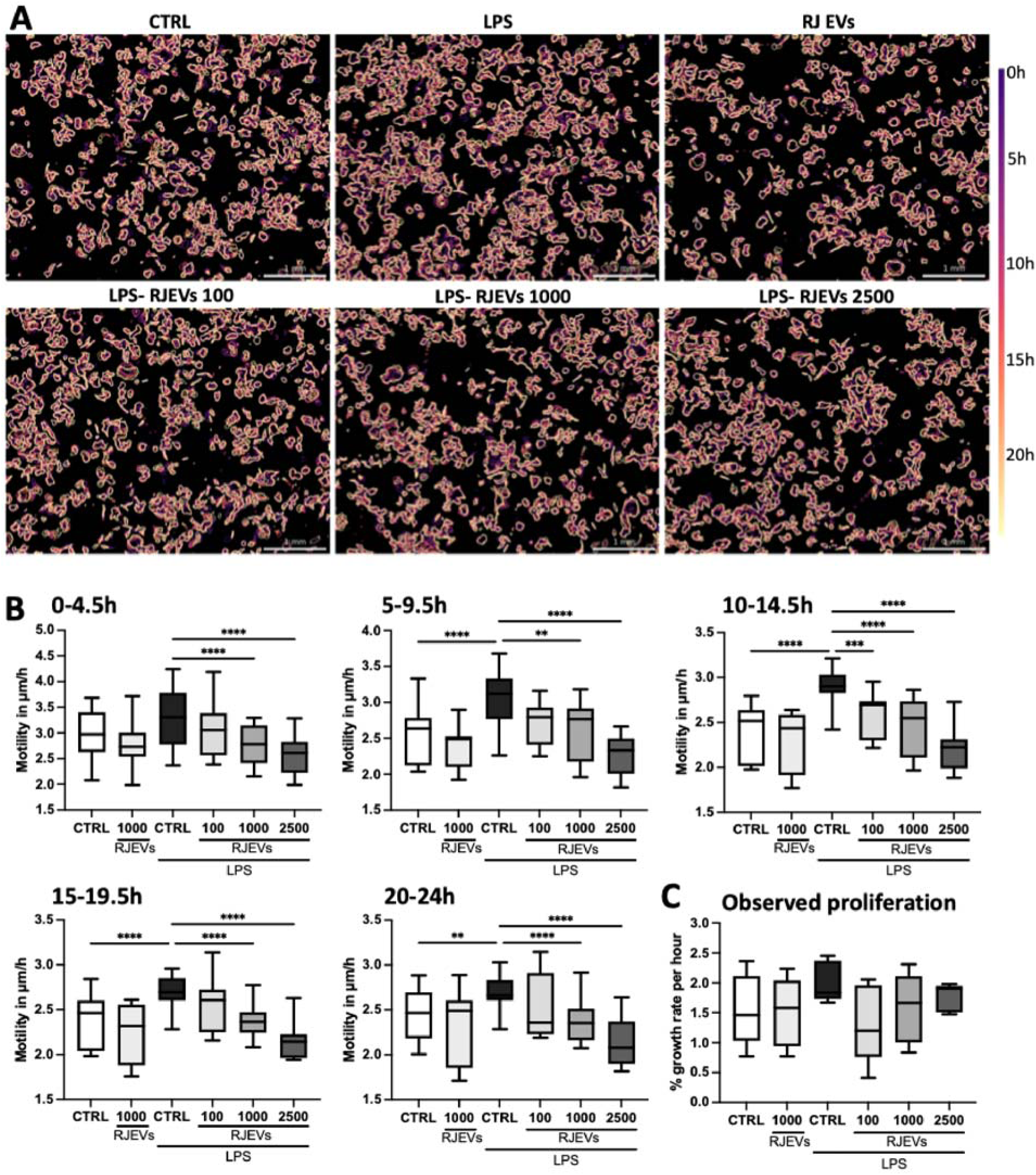
Effect of LPS and RJEVs on microglial motility; Cellular motility was measured with a PHIO cell watcher M; images were taken automatically every 30min; A) Cell map displaying cellular outlines for HMC3 cells in each condition over the time course of 24h, with purple representing 0h and yellow 24h. B) Motility of HMC3 cells in µm/h for different phases of exposure to LPS and RJEVs: 0-4.5h, 5-9.5h, 10-14.5h, 15-19.5h, 20-24h; C) Proliferation observed with PHIO cell watcher, displayed as percentage growth rate per hour; data is displayed as box and whiskers (min-max); n=5; Significance was tested with Kruskal-Wallis test and Dunn’s multiple comparison; *p<0.05, **p<0.01, ***p<0.001, ****p<0.0001;

### Modulation of cell elasticity by RJEVs depends on the presence of LPS

The Young’s modulus-a measure for cell stiffness and elasticity-of HMC3 cells in different conditions was assessed with AFM. **Figure 5A** shows representative nanomechanical force maps for scan height and elasticity for each group. The overall cell elasticity is displayed as all pooled force curves of a minimum of 4 independent experiments per group (**Figure 5B**). The control group (2412 values) displayed a median Young’s modulus of 4205 Pa (2817 Pa 25% Percentile, 6486 Pa 75% Percentile). Incubation with RJEV or LPS decreased stiffness by 19.25% and 54.34% respectively, compared to the untreated control (RJEV median 3395 Pa, 2485 Pa 25% Percentile, 3395 Pa 75% Percentile; LPS median 1920 Pa, 1412 Pa 25% Percentile, 4648 Pa 75% Percentile). Interestingly, treatment with RJEV and LPS resulted in the opposite effect and significantly increased median cell stiffness by 76.6% compared to untreated control (**Figure 5 B and 5C**).

**Figure 5:**
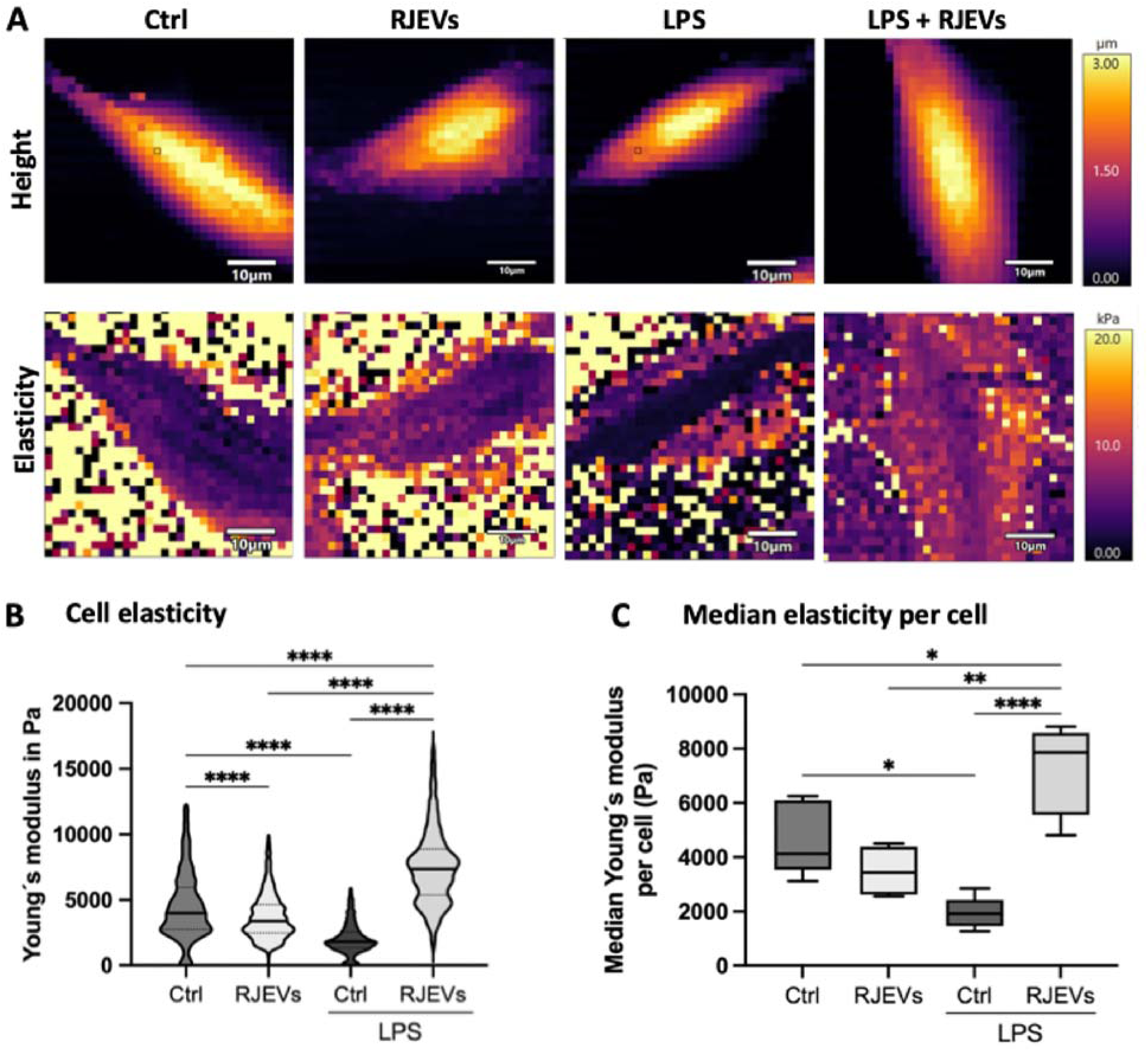
Impact of LPS and RJEVs on microglial nano-mechanics as probed by atomic force microscopy (AFM). (A) Height and elasticity (Pa) force maps of living microglial cells in buffer, where each pixel represents a force-distance curve from where indentation data was obtained. (B) Violin plot of pooled Young’s modulus (in Pa) for all force curves obtained on control and RJEV, LPS, and LPS-RJEV exposed cells. (C) Overall cell Young’s modulus medians (in Pa) across the four experimental conditions, with and without LPS exposure. Significance was tested with Kruskal-Wallis test and Dunn’s multiple comparison; *p<0.05, **p<0.01, ****p<0.0001.

### Assessing membrane fluidity with FRAP

Molecular dynamics of the HMC3 cell membrane in different conditions were measured with FRAP. **Figure 6A** shows representative cell images and photobleached areas before and after exposure to the laser. Quantitatively, the half-recovery time t_1/2_, rate constant K1, and mobile fraction were determined. Control cells displayed a median t_1/2_ of 2.78 s (25% percentile 2.13 s, 75% percentile 4.03 s). RJEVs significantly decreased half recovery time (3.1 s, 25% percentile 2.46 s, 75% percentile 3.93 s), while LPS significantly increased it (2.17 s, 25% percentile 1.55 s, 75% percentile 2.8 s). RJEVs increased the half recovery time of LPS-stimulated cells (2.5 s, 25% percentile 1.84 s, 75% percentile 3.11 s) (**Figure 6Bi**). The rate constant was not significantly changed by RJEVs (ctrl: 0.22 µm^2^/s, 25% percentile 0.15 µm^2^/s, 75% percentile 0.32 µm^2^/s; RJEVs 0.22 µm^2^/s, 25% percentile 0.18 µm^2^/s, 75% percentile 0.28 µm^2^/s). However, LPS stimulation significantly increased the rate constant (0.32 µm^2^/s, 25% percentile 0.25 µm^2^/s, 75% percentile 0.45 µm^2^/s). Treatment of LPS-stimulated cells with RJEVs significantly decreased the rate constant, close to levels of unstimulated cells (0.25 µm^2^/s, 25% percentile 0.20 µm^2^/s, 75% percentile 0.37 µm^2^/s) (**Figure 6Bii**). Interestingly, the mobile fraction was similar between groups (77.2%-82%) and did not display significant differences (**Figure 6Biii**).

**Figure 6:**
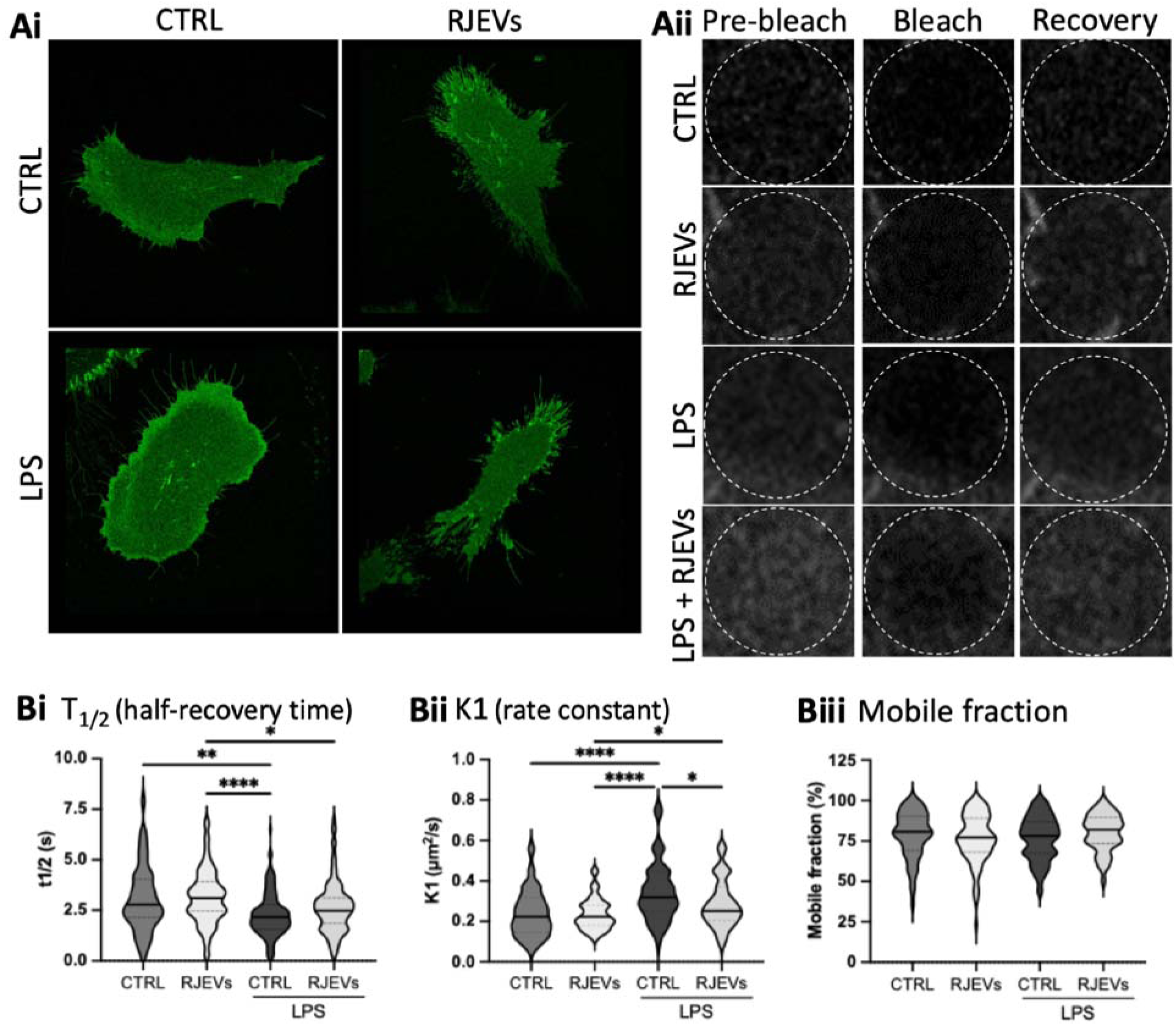
Fluorescence Recovery After Photobleaching (FRAP); Ai) representative images of HMC3 cells stained with Cell mask; Aii) magnification of photobleached area before (pre-bleach), during (bleach) and after photobleaching (recovery); B) Quantification of fluorescence recovery of untreated control cells (ctrl) and cells treated with RJEVs, LPS, or both; Bi) T1/2 half-recovery time, Bii) K1 rate constant and Biii) Mobile fraction; n=3-5 different coverslips and 84-122 separate measurements. Data is displayed as violin plot with median (line) and quartiles (dotted lines). Significance was tested with Kruskal-Wallis test and Dunn’s multiple comparison; *p<0.05, **p<0.01, ***p<0.001, ****p<0.0001;

## DISCUSSION

Neuroinflammation is recognized as one of the hallmarks of numerous neurodegenerative and psychiatric diseases. A recent perspective paper highlighted the importance of multidisciplinary approaches to unravel underlying mechanisms and design novel treatment options for these conditions (23). Molecular pathways have been studied in detail, however, mechanobiological responses of key cellular players reacting to relevant pro-inflammatory stimuli are widely understudied. It is generally understood that cells require outer stimuli to activate or trigger mechano-sensitive receptors or effects. However, one of the most underestimated mechanical stimuli is the cell membrane itself, not only in pathological changes but also as a target for novel treatments. Thus, in this study we used a multi-disciplinary approach to analyze the activation of human microglial cells by LPS and the subsequent treatment with RJEVs.

As a first step, we assessed the effect of LPS and RJEV on microglial cell viability (MTT-, LDH assay) and proliferation (**Figure 2A, B, C**). As reported by other groups (24,25), LPS was found to increase cell proliferation, but this effect was counteracted by RJEVs in a dose-dependent manner. Interestingly, RJEVs reverted the negative effects of LPS on cell viability, but did not display the same dose-dependent effect for cell viability, indicating a therapeutic window rather than a linear effect. While 10-2500 RJEV/cell increased metabolic activity, 1 and 5000 JREV/cell had no significant effect, reason for which the higher concentration was excluded from further testing.

It has been previously described that bulk RJ displayed anti-inflammatory properties in microglia by You et al (26); however, the effect of isolated RJEVs remained unknown. In this study, we are the first to describe the anti-inflammatory effect of RJEVs in microglia in a dose-dependent manner for all three tested cytokines (IL1b, IL6 and TNFα) (**Figure 2D**). Interestingly, in this assay, we observed a saturation point at 1000 RJEV/cell and no further decrease in cytokine secretion at 2500 RJEVs. A similar effect has been described by Malvicini et al (27) in LPS-stimulated macrophages treated with mesenchymal stromal cell-derived small EVs: while the anti-inflammatory effect increased from 0.05-0.5×10^10^ EVs, there was no additional beneficial effect observed for 1×10^10^. We hypothesize that due to the endpoint testing with ELISA, subtle differences are being lost, as indeed, significant differences between 1000 RJEVs and 2500 RJEVs per cell were observed in the PHIO motility assay between 5 and 14.5 hours (**Supplemental Table 3**).

There are copious studies on the effects of different EVs on microglia (28–30), but so far little emphasis has been placed on how these EVs exert their biological effects upon entering the cell. It is widely accepted that EVs are being internalized into target cells, but a crucial factor is the route of uptake that determines their fate within the cell (summarized by O’Brien et al. (31)). So far, no studies have investigated the routes of internalization of EVs into microglia, especially comparing differences between activated and resting state cells. We found significant differences in RJEV uptake between the two activation states: not only did LPS-stimulated, activated microglia internalize significantly more RJEVs, but there was also a distinct shift in the distribution of internalization routes (**Figure 3**). In the resting state, an equilibrium between the four mechanisms can be observed, with all routes being involved in RJEV uptake. However, upon LPS stimulation, lipid-raft mediated uptake is being reduced from 21% to around 4% of total RJEVs internalized. Miller et al. described a pronounced change in lipid rafts in inflammatory states, referring to them as “inflammarafts”, which are likely not favoring RJEV uptake (32). At the same time, RJEV uptake via clathrin-dependent endocytosis and macropinocytosis significantly increased following LPS exposure. Both mechanisms are highly dependent on actin organization, a hallmark of mechanobiology (33,34). These results suggest that activated microglia actively alter their preferred EV internalization routes compared to resting state cells, which could potentially be exploited as a novel alternative for the targeted treatment of neuroinflammation.

Microglia are an active part of the central nervous system (CNS) immune system, continuously migrating and scavenging to maintain tissue homeostasis (35). However, after sustained activation by pro-inflammatory stimuli such as LPS, microglia not only increase their secretion of pro-inflammatory cytokines but also their motility (36), which again can lead to detrimental effects. We tracked the motility of HMC3 microglia with a PHIO cell watcher for 24 hours (**Figure 4**) and observed, indeed, an increase in motility after stimulation with LPS. RJEVs decreased motility below the control baseline level. However, this effect was specific for the presence of LPS, as in control conditions with RJEVs, no differences were noted. This is an effect that so far has not been reported, neither for RJ nor for other compounds. However, it is known that activation of microglia leads to a dramatic reorganization of their cytoskeleton (37), which could be stabilized by RJEVs, leading to more pronounced effects. Even more so, as described above, there is a significant change in both the route of uptake and amount of RJEVs internalized.

To further understand the nanomechanical impact of LPS and RJEVs exposure on microglia, force-mapping of living HMC3 cells was performed with AFM (22). Firstly, we observed that Young’s modulus for living HMC3 cells is in the low kPa range which is consistent with the expected elasticity range for glial cells in buffer (38). Interestingly, exposure to LPS as well as to RJEVs reduced cellular stiffness; however, when LPS and RJEVs were combined, stiffness significantly increased (**Figure 5**). Alterations in the biological behavior of cells are associated with changes in cellular mechanics in a bidirectional manner (39). Previous studies have shed light on the role of LPS and inflammation on the mechanical properties of cell membranes with contradictory results. For example, LPS exposure has been seen to increase the stiffness of macrophages and endothelial cells (40,41). However, similar to our current results, a decrease in cell elasticity has been observed in lung epithelial cells (42) and white matter-derived microglia (43), following incubation with LPS. Several potential mechanisms can explain these changes such as reorganization of the cell cytoskeleton (44–46), changes in membrane fluidity (47), and the direct effect of inflammatory cytokines on cellular reorganization (48). More importantly, the correlation between our AFM data and FRAP findings (**Figure 6**) suggests that altered membrane fluidity is one of the main drivers behind changes in single-cell mechanics of glial cells during neuroinflammation, which can be reverted with RJEV treatment. The anti-inflammatory effect of RJEVs further suggests that this reduction in Young’s modulus following LPS exposure is a mechanobiological hallmark of HMC3 inflammation and can explain why RJEV treatment counteracts these nanomechanical changes. Interestingly, other previous work has shown that triptolide, a diterpenoid epoxide suspected to affect membrane fluidity, also possesses anti-inflammatory effects that correlated with increased cell stiffness (49). Nevertheless, future work should investigate the potential mechanisms behind these observations.

It is well known that exposure to environmental factors such as inflammatory mediators and anti-inflammatory molecules can modulate the biological behavior of neural cells, among others (50,51). What is less known, is how these changes are associated with alterations in cellular mechanical properties such as elasticity and fluidity. Here we show that there is an important correlation between cell stiffness, membrane fluidity, and motility and migration in HMC3 cells, strongly suggesting a direct interplay between nanomechanical properties and physiological behavior. Particularly, we observed that LPS-mediated inflammation leads to reduced cell Young’s modulus but increased membrane fluidity, which in turn was associated to increased migration and motility (**Figure 4 and 5**). This data suggests that the reorganization of both the cell membrane and cytoskeleton is crucial for microglial motility and plays a crucial role in its biological response during neuroinflammation.

## CONCLUSION

This study provides the first detailed mechanistic analysis of inflammation-associated mechanobiological changes in microglia, and how a novel EV-based therapy based on RJEVs can modulate these alterations. We demonstrate that LPS activation significantly alters microglial biomechanics by decreasing Young’s modulus and increasing membrane fluidity and motility—changes that are effectively counteracted by RJEV treatment. Furthermore, our findings establish a link between microglial activation, vesicle uptake mechanisms, and changes in cellular nanomechanics, strongly suggesting that alterations in cell mechanobiology are an integral factor in the development of neuroinflammation. Furthermore, these results expand our understanding of how EVs interact with microglia, demonstrating that their biological effects extend beyond biochemical signaling to include modulation of cellular mechanics and membrane fluidity. Given the increasing importance of mechanobiology in health and disease, these findings add another therapeutic layer to the potential of EV-based therapies for neuroinflammation and beyond. We expect that these findings will pave the way towards the understanding of the interplay between microglial biology and mechanical properties, as a potential key target for preventing and treating neuroinflammation in the future.

## DECLARATIONS

### AUTHOR CONTRIBUTION

**GZ and PB:** Conceptualization, data curation, investigation, formal analysis, methodology, visualization, writing-original draft, review and editing. **FS, GC, JAM**: Data curation, investigation and writing-review and editing. **NB, PDC:** Investigation, formal analysis, supervision, validation, discussion. **NB:** funding acquisition. **SA:** Methodology, investigation, formal analysis, supervision, validation, funding acquisition, writing-original draft, review and editing. **CMAPS:** Conceptualization, methodology, supervision, validation, project administration, formal analysis, data curation, funding acquisition, validation, writing-original draft, review and editing; All authors approved the final version of the manuscript.

## ACKNOWLEDGEMENTS

The authors would like to thank the Advanced Microscopy Unit (UMA UC) for their support with the TEM. The graphical abstract was created with BioRender, license for publication was purchased by CMAPS.

