## Supplemental Material for "Royal Jelly Derived Extracellular Vesicles Modulate Microglial Nanomechanics and Inflammatory Responses"

**Supplemental table 1**: Descriptive statistics figure 2

| **TNF alpha** | ctrl | RJEV 1000/c | LPS | LPS Indom. | LPS RJEV 1/c | LPS RJEV 10/c | LPS RJEV 100/c | LPS RJEV 1000 | LPS RJEV 2500/c |
| --- | --- | --- | --- | --- | --- | --- | --- | --- | --- |
| Minimum | 2.1 | 2.1 | 487 | 81.69 | 495 | 453 | 270.2 | 140.3 | 134.1 |
| Maximum | 5.489 | 4.466 | 554.3 | 128 | 558.3 | 483.1 | 316 | 192 | 187 |
| Mean | 3.559 | 3.059 | 519.6 | 102.8 | 519.9 | 470.9 | 287.1 | 166.2 | 161.1 |
| Std. Deviation | 1.582 | 1.018 | 30.29 | 19.63 | 29.37 | 13.88 | 21.2 | 21.23 | 21.69 |
| **IL-1** | ctrl | RJEV 1000/c | LPS | LPS Indom. | LPS RJEV 1/c | LPS RJEV 10/c | LPS RJEV 100/c | LPS RJEV 1000 | LPS RJEV 2500/c |
| Minimum | 5 | 6 | 338 | 64 | 341 | 333 | 255 | 125 | 130 |
| Maximum | 13 | 12 | 389 | 91 | 393 | 371 | 292 | 171 | 170 |
| Mean | 8.25 | 9 | 362 | 79.25 | 360.5 | 348 | 277.3 | 151.3 | 152.5 |
| Std. Deviation | 3.594 | 2.582 | 22.29 | 12.97 | 24.24 | 17.78 | 16.72 | 20.66 | 17.08 |
| **IL-6** | ctrl | RJEV 1000/c | LPS | LPS Indom. | LPS RJEV 1/c | LPS RJEV 10/c | LPS RJEV 100/c | LPS RJEV 1000 | LPS RJEV 2500/c |
| Minimum | 6.34 | 7.09 | 188.9 | 25.55 | 183.4 | 164.7 | 131.1 | 111 | 103.8 |
| Maximum | 19.46 | 15.68 | 266.7 | 55.69 | 254.6 | 249 | 189.5 | 160.7 | 140.8 |
| Mean | 12.23 | 11.21 | 230.1 | 36.76 | 223.7 | 212.9 | 159.7 | 135.3 | 123.6 |
| Std. Deviation | 5.674 | 4.531 | 32.2 | 13.1 | 30.22 | 35.73 | 23.89 | 20.55 | 16.09 |

**Supplemental table 1**: Statistical analysis figure 2

| **TNF alpha** |  |  |  |  |
| --- | --- | --- | --- | --- |
| **Tukey's multiple comparisons test** | **Mean Diff.** | **95.00% CI of diff.** | **Summary** | **Adj. P Value** |
| ctrl vs. RJEV 1000/c | 0.4998 | -47.95 to 48.95 | ns | >0.9999 |
| ctrl vs. LPS | -516 | -564.5 to -467.6 | **** | <0.0001 |
| ctrl vs. LPS Indom. | -99.26 | -147.7 to -50.81 | **** | <0.0001 |
| ctrl vs. LPS RJEV 1/c | -516.3 | -564.8 to -467.9 | **** | <0.0001 |
| ctrl vs. LPS RJEV 10/c | -467.4 | -515.8 to -418.9 | **** | <0.0001 |
| ctrl vs. LPS RJEV 100/c | -283.6 | -332.0 to -235.1 | **** | <0.0001 |
| ctrl vs. LPS RJEV 1000 | -162.6 | -211.1 to -114.2 | **** | <0.0001 |
| ctrl vs. LPS RJEV 2500/c | -157.5 | -206.0 to -109.1 | **** | <0.0001 |
| RJEV 1000/c vs. LPS | -516.5 | -565.0 to -468.1 | **** | <0.0001 |
| RJEV 1000/c vs. LPS Indom. | -99.76 | -148.2 to -51.31 | **** | <0.0001 |
| RJEV 1000/c vs. LPS RJEV 1/c | -516.8 | -565.3 to -468.4 | **** | <0.0001 |
| RJEV 1000/c vs. LPS RJEV 10/c | -467.9 | -516.3 to -419.4 | **** | <0.0001 |
| RJEV 1000/c vs. LPS RJEV 100/c | -284.1 | -332.5 to -235.6 | **** | <0.0001 |
| RJEV 1000/c vs. LPS RJEV 1000 | -163.1 | -211.6 to -114.7 | **** | <0.0001 |
| RJEV 1000/c vs. LPS RJEV 2500/c | -158 | -206.5 to -109.6 | **** | <0.0001 |
| LPS vs. LPS Indom. | 416.8 | 368.3 to 465.2 | **** | <0.0001 |
| LPS vs. LPS RJEV 1/c | -0.3224 | -48.77 to 48.13 | ns | >0.9999 |
| LPS vs. LPS RJEV 10/c | 48.66 | 0.2133 to 97.11 | * | 0.0484 |
| LPS vs. LPS RJEV 100/c | 232.5 | 184.0 to 280.9 | **** | <0.0001 |
| LPS vs. LPS RJEV 1000 | 353.4 | 304.9 to 401.8 | **** | <0.0001 |
| LPS vs. LPS RJEV 2500/c | 358.5 | 310.0 to 406.9 | **** | <0.0001 |
| LPS Indom. vs. LPS RJEV 1/c | -417.1 | -465.5 to -368.6 | **** | <0.0001 |
| LPS Indom. vs. LPS RJEV 10/c | -368.1 | -416.5 to -319.6 | **** | <0.0001 |
| LPS Indom. vs. LPS RJEV 100/c | -184.3 | -232.8 to -135.9 | **** | <0.0001 |
| LPS Indom. vs. LPS RJEV 1000 | -63.37 | -111.8 to -14.92 | ** | 0.0041 |
| LPS Indom. vs. LPS RJEV 2500/c | -58.27 | -106.7 to -9.821 | ** | 0.01 |
| LPS RJEV 1/c vs. LPS RJEV 10/c | 48.99 | 0.5357 to 97.44 | * | 0.046 |
| LPS RJEV 1/c vs. LPS RJEV 100/c | 232.8 | 184.3 to 281.2 | **** | <0.0001 |
| LPS RJEV 1/c vs. LPS RJEV 1000 | 353.7 | 305.3 to 402.2 | **** | <0.0001 |
| LPS RJEV 1/c vs. LPS RJEV 2500/c | 358.8 | 310.4 to 407.3 | **** | <0.0001 |
| LPS RJEV 10/c vs. LPS RJEV 100/c | 183.8 | 135.3 to 232.2 | **** | <0.0001 |
| LPS RJEV 10/c vs. LPS RJEV 1000 | 304.7 | 256.3 to 353.2 | **** | <0.0001 |
| LPS RJEV 10/c vs. LPS RJEV 2500/c | 309.8 | 261.4 to 358.3 | **** | <0.0001 |
| LPS RJEV 100/c vs. LPS RJEV 1000 | 120.9 | 72.48 to 169.4 | **** | <0.0001 |
| LPS RJEV 100/c vs. LPS RJEV 2500/c | 126 | 77.58 to 174.5 | **** | <0.0001 |
| LPS RJEV 1000 vs. LPS RJEV 2500/c | 5.103 | -43.35 to 53.55 | ns | >0.9999 |

**TNF alpha**

| **Tukey's multiple comparisons test** | **Mean Diff.** | **95.00% CI of diff.** | **Summary** | **Adj. P Value** |
| --- | --- | --- | --- | --- |
| ctrl vs. RJEV 1000/c | -0.75 | -41.08 to 39.58 | ns | >0.9999 |
| ctrl vs. LPS | -353.8 | -394.1 to -313.4 | **** | <0.0001 |
| ctrl vs. LPS Indom. | -71 | -111.3 to -30.67 | **** | <0.0001 |
| ctrl vs. LPS RJEV 1/c | -352.3 | -392.6 to -311.9 | **** | <0.0001 |
| ctrl vs. LPS RJEV 10/c | -339.8 | -380.1 to -299.4 | **** | <0.0001 |
| ctrl vs. LPS RJEV 100/c | -269 | -309.3 to -228.7 | **** | <0.0001 |
| ctrl vs. LPS RJEV 1000 | -143 | -183.3 to -102.7 | **** | <0.0001 |
| ctrl vs. LPS RJEV 2500/c | -144.3 | -184.6 to -103.9 | **** | <0.0001 |
| RJEV 1000/c vs. LPS | -353 | -393.3 to -312.7 | **** | <0.0001 |
| RJEV 1000/c vs. LPS Indom. | -70.25 | -110.6 to -29.92 | **** | <0.0001 |
| RJEV 1000/c vs. LPS RJEV 1/c | -351.5 | -391.8 to -311.2 | **** | <0.0001 |
| RJEV 1000/c vs. LPS RJEV 10/c | -339 | -379.3 to -298.7 | **** | <0.0001 |
| RJEV 1000/c vs. LPS RJEV 100/c | -268.3 | -308.6 to -227.9 | **** | <0.0001 |
| RJEV 1000/c vs. LPS RJEV 1000 | -142.3 | -182.6 to -101.9 | **** | <0.0001 |
| RJEV 1000/c vs. LPS RJEV 2500/c | -143.5 | -183.8 to -103.2 | **** | <0.0001 |
| LPS vs. LPS Indom. | 282.8 | 242.4 to 323.1 | **** | <0.0001 |
| LPS vs. LPS RJEV 1/c | 1.5 | -38.83 to 41.83 | ns | >0.9999 |
| LPS vs. LPS RJEV 10/c | 14 | -26.33 to 54.33 | ns | 0.9566 |
| LPS vs. LPS RJEV 100/c | 84.75 | 44.42 to 125.1 | **** | <0.0001 |
| LPS vs. LPS RJEV 1000 | 210.8 | 170.4 to 251.1 | **** | <0.0001 |
| LPS vs. LPS RJEV 2500/c | 209.5 | 169.2 to 249.8 | **** | <0.0001 |
| LPS Indom. vs. LPS RJEV 1/c | -281.3 | -321.6 to -240.9 | **** | <0.0001 |
| LPS Indom. vs. LPS RJEV 10/c | -268.8 | -309.1 to -228.4 | **** | <0.0001 |
| LPS Indom. vs. LPS RJEV 100/c | -198 | -238.3 to -157.7 | **** | <0.0001 |
| LPS Indom. vs. LPS RJEV 1000 | -72 | -112.3 to -31.67 | **** | <0.0001 |
| LPS Indom. vs. LPS RJEV 2500/c | -73.25 | -113.6 to -32.92 | **** | <0.0001 |
| LPS RJEV 1/c vs. LPS RJEV 10/c | 12.5 | -27.83 to 52.83 | ns | 0.9776 |
| LPS RJEV 1/c vs. LPS RJEV 100/c | 83.25 | 42.92 to 123.6 | **** | <0.0001 |
| LPS RJEV 1/c vs. LPS RJEV 1000 | 209.3 | 168.9 to 249.6 | **** | <0.0001 |
| LPS RJEV 1/c vs. LPS RJEV 2500/c | 208 | 167.7 to 248.3 | **** | <0.0001 |
| LPS RJEV 10/c vs. LPS RJEV 100/c | 70.75 | 30.42 to 111.1 | **** | <0.0001 |
| LPS RJEV 10/c vs. LPS RJEV 1000 | 196.8 | 156.4 to 237.1 | **** | <0.0001 |
| LPS RJEV 10/c vs. LPS RJEV 2500/c | 195.5 | 155.2 to 235.8 | **** | <0.0001 |
| LPS RJEV 100/c vs. LPS RJEV 1000 | 126 | 85.67 to 166.3 | **** | <0.0001 |
| LPS RJEV 100/c vs. LPS RJEV 2500/c | 124.8 | 84.42 to 165.1 | **** | <0.0001 |
| LPS RJEV 1000 vs. LPS RJEV 2500/c | -1.25 | -41.58 to 39.08 | ns | >0.9999 |

| **IL-6** |  |  |  |  |
| --- | --- | --- | --- | --- |
| **Tukey's multiple comparisons test** | **Mean Diff.** | **95.00% CI of diff.** | **Summary** | **Adj. P Value** |
| ctrl vs. RJEV 1000/c | 1.019 | -53.37 to 55.41 | ns | >0.9999 |
| ctrl vs. LPS | -217.8 | -272.2 to -163.4 | **** | <0.0001 |
| ctrl vs. LPS Indom. | -24.53 | -78.92 to 29.85 | ns | 0.8376 |
| ctrl vs. LPS RJEV 1/c | -211.5 | -265.9 to -157.1 | **** | <0.0001 |
| ctrl vs. LPS RJEV 10/c | -200.7 | -255.1 to -146.3 | **** | <0.0001 |
| ctrl vs. LPS RJEV 100/c | -147.4 | -201.8 to -93.06 | **** | <0.0001 |
| ctrl vs. LPS RJEV 1000 | -123 | -177.4 to -68.64 | **** | <0.0001 |
| ctrl vs. LPS RJEV 2500/c | -111.4 | -165.8 to -56.99 | **** | <0.0001 |
| RJEV 1000/c vs. LPS | -218.8 | -273.2 to -164.5 | **** | <0.0001 |
| RJEV 1000/c vs. LPS Indom. | -25.55 | -79.94 to 28.83 | ns | 0.8065 |
| RJEV 1000/c vs. LPS RJEV 1/c | -212.5 | -266.9 to -158.1 | **** | <0.0001 |
| RJEV 1000/c vs. LPS RJEV 10/c | -201.7 | -256.1 to -147.3 | **** | <0.0001 |
| RJEV 1000/c vs. LPS RJEV 100/c | -148.5 | -202.9 to -94.08 | **** | <0.0001 |
| RJEV 1000/c vs. LPS RJEV 1000 | -124 | -178.4 to -69.66 | **** | <0.0001 |
| RJEV 1000/c vs. LPS RJEV 2500/c | -112.4 | -166.8 to -58.01 | **** | <0.0001 |
| LPS vs. LPS Indom. | 193.3 | 138.9 to 247.7 | **** | <0.0001 |
| LPS vs. LPS RJEV 1/c | 6.34 | -48.05 to 60.73 | ns | >0.9999 |
| LPS vs. LPS RJEV 10/c | 17.14 | -37.25 to 71.53 | ns | 0.9752 |
| LPS vs. LPS RJEV 100/c | 70.38 | 15.99 to 124.8 | ** | 0.0046 |
| LPS vs. LPS RJEV 1000 | 94.79 | 40.41 to 149.2 | **** | <0.0001 |
| LPS vs. LPS RJEV 2500/c | 106.4 | 52.06 to 160.8 | **** | <0.0001 |
| LPS Indom. vs. LPS RJEV 1/c | -186.9 | -241.3 to -132.6 | **** | <0.0001 |
| LPS Indom. vs. LPS RJEV 10/c | -176.1 | -230.5 to -121.8 | **** | <0.0001 |
| LPS Indom. vs. LPS RJEV 100/c | -122.9 | -177.3 to -68.53 | **** | <0.0001 |
| LPS Indom. vs. LPS RJEV 1000 | -98.5 | -152.9 to -44.11 | **** | <0.0001 |
| LPS Indom. vs. LPS RJEV 2500/c | -86.84 | -141.2 to -32.46 | *** | 0.0003 |
| LPS RJEV 1/c vs. LPS RJEV 10/c | 10.8 | -43.59 to 65.19 | ns | 0.9988 |
| LPS RJEV 1/c vs. LPS RJEV 100/c | 64.04 | 9.649 to 118.4 | * | 0.0123 |
| LPS RJEV 1/c vs. LPS RJEV 1000 | 88.45 | 34.07 to 142.8 | *** | 0.0003 |
| LPS RJEV 1/c vs. LPS RJEV 2500/c | 100.1 | 45.72 to 154.5 | **** | <0.0001 |
| LPS RJEV 10/c vs. LPS RJEV 100/c | 53.24 | -1.151 to 107.6 | ns | 0.0585 |
| LPS RJEV 10/c vs. LPS RJEV 1000 | 77.65 | 23.27 to 132.0 | ** | 0.0015 |
| LPS RJEV 10/c vs. LPS RJEV 2500/c | 89.31 | 34.92 to 143.7 | *** | 0.0002 |
| LPS RJEV 100/c vs. LPS RJEV 1000 | 24.42 | -29.97 to 78.80 | ns | 0.841 |
| LPS RJEV 100/c vs. LPS RJEV 2500/c | 36.07 | -18.32 to 90.46 | ns | 0.4149 |
| LPS RJEV 1000 vs. LPS RJEV 2500/c | 11.65 | -42.73 to 66.04 | ns | 0.998 |

**Supplemental table 3**: Descriptive statistics figure 4

| **0 to 4.5h** | **ctrl** | **LPS** | **RJEVs** | **LPS RJEVs 100** | **LPS RJEVs 1000** | **LPS RJEVs 2500** |
| --- | --- | --- | --- | --- | --- | --- |
| Minimum | 2.077 | 2.363 | 1.987 | 2.381 | 2.156 | 1.988 |
| Maximum | 3.683 | 4.243 | 3.719 | 4.187 | 3.294 | 3.287 |
| Mean | 2.946 | 3.29 | 2.768 | 3.054 | 2.75 | 2.562 |
| Std. Deviation | 0.4634 | 0.5429 | 0.3501 | 0.4828 | 0.3684 | 0.3733 |
| **5 to 9.5h** | **ctrl** | **LPS** | **RJEVs** | **LPS RJEVs 100** | **LPS RJEVs 1000** | **LPS RJEVs 2500** |
| Minimum | 2.036 | 2.262 | 1.921 | 2.251 | 1.96 | 1.817 |
| Maximum | 3.33 | 3.677 | 2.894 | 3.16 | 3.181 | 2.664 |
| Mean | 2.535 | 2.983 | 2.35 | 2.687 | 2.594 | 2.266 |
| Std. Deviation | 0.3841 | 0.4242 | 0.255 | 0.2846 | 0.4091 | 0.28 |
| **10 to 14.5h** | **ctrl** | **LPS** | **RJEVs** | **LPS RJEVs 100** | **LPS RJEVs 1000** | **LPS RJEVs 2500** |
| Minimum | 1.975 | 2.419 | 1.771 | 2.215 | 1.965 | 1.883 |
| Maximum | 2.796 | 3.211 | 2.638 | 2.954 | 2.862 | 2.728 |
| Mean | 2.387 | 2.898 | 2.271 | 2.586 | 2.447 | 2.233 |
| Std. Deviation | 0.3223 | 0.1909 | 0.3362 | 0.2647 | 0.3128 | 0.2733 |
| **15 to 19.5h** | **ctrl** | **LPS** | **RJEVs** | **LPS RJEVs 100** | **LPS RJEVs 1000** | **LPS RJEVs 2500** |
| Minimum | 1.985 | 2.281 | 1.757 | 2.158 | 2.08 | 1.943 |
| Maximum | 2.841 | 2.96 | 2.611 | 3.14 | 2.775 | 2.627 |
| Mean | 2.37 | 2.692 | 2.222 | 2.54 | 2.403 | 2.168 |
| Std. Deviation | 0.3036 | 0.1951 | 0.3354 | 0.295 | 0.2032 | 0.2314 |
| **15 to 19.5h** | **ctrl** | **LPS** | **RJEVs** | **LPS RJEVs 100** | **LPS RJEVs 1000** | **LPS RJEVs 2500** |
| Minimum | 2.005 | 2.284 | 1.711 | 2.191 | 2.072 | 1.816 |
| Maximum | 2.885 | 3.03 | 2.888 | 3.148 | 2.915 | 2.637 |
| Mean | 2.447 | 2.687 | 2.264 | 2.556 | 2.39 | 2.163 |
| Std. Deviation | 0.2911 | 0.1968 | 0.3997 | 0.3616 | 0.2533 | 0.284 |
| **Growth rate** | **ctrl** | **LPS** | **RJEVs** | **LPS RJEVs 100** | **LPS RJEVs 1000** | **LPS RJEVs 2500** |
| Minimum | 0.7748 | 1.672 | 0.7733 | 0.4146 | 0.8364 | 1.475 |
| Maximum | 2.362 | 2.454 | 2.238 | 2.057 | 2.31 | 1.985 |
| Mean | 1.551 | 2.006 | 1.509 | 1.329 | 1.581 | 1.764 |
| Std. Deviation | 0.599 | 0.3413 | 0.5806 | 0.6554 | 0.5848 | 0.2407 |

**Supplemental table 4:** Statistical analysis motility

| **0 to 4.5 hours** |  |  |  |  |
| --- | --- | --- | --- | --- |
| **Dunn's multiple comparisons test** | **Mean rank diff.** | **Significant?** | **Summary** | **Adjusted P Value** |
| ctrl vs. LPS | -40.48 | No | ns | 0.2149 |
| ctrl vs. RJ | 31.2 | No | ns | 0.8866 |
| ctrl vs. LPS RJEV 100 | -13.15 | No | ns | >0.9999 |
| ctrl vs. LPS RJEV 1000 | 33.16 | No | ns | 0.6723 |
| ctrl vs. LPS RJEV 2500 | 64.82 | Yes | ** | 0.0013 |
| LPS vs. RJ | 71.68 | Yes | *** | 0.0002 |
| LPS vs. LPS RJEV 100 | 27.33 | No | ns | >0.9999 |
| LPS vs. LPS RJEV 1000 | 73.64 | Yes | **** | <0.0001 |
| LPS vs. LPS RJEV 2500 | 105.3 | Yes | **** | <0.0001 |
| RJ vs. LPS RJEV 100 | -44.34 | No | ns | 0.0998 |
| RJ vs. LPS RJEV 1000 | 1.967 | No | ns | >0.9999 |
| RJ vs. LPS RJEV 2500 | 33.62 | No | ns | 0.5944 |
| LPS RJEV 100 vs. LPS RJEV 1000 | 46.31 | No | ns | 0.0689 |
| LPS RJEV 100 vs. LPS RJEV 2500 | 77.97 | Yes | **** | <0.0001 |
| LPS RJEV 1000 vs. LPS RJEV 2500 | 31.66 | No | ns | 0.7907 |
| **5 to 9.5 hours** |  |  |  |  |
| **Dunn's multiple comparisons test** | **Mean rank diff.** | **Significant?** | **Summary** | **Adjusted P Value** |
| ctrl vs. LPS | -84.29 | Yes | **** | <0.0001 |
| ctrl vs. RJ | 39.6 | No | ns | 0.3573 |
| ctrl vs. LPS RJEV 100 | -38.42 | No | ns | 0.4248 |
| ctrl vs. LPS RJEV 1000 | -15.45 | No | ns | >0.9999 |
| ctrl vs. LPS RJEV 2500 | 58.51 | Yes | * | 0.0126 |
| LPS vs. RJ | 123.9 | Yes | **** | <0.0001 |
| LPS vs. LPS RJEV 100 | 45.87 | No | ns | 0.1326 |
| LPS vs. LPS RJEV 1000 | 68.84 | Yes | ** | 0.0013 |
| LPS vs. LPS RJEV 2500 | 142.8 | Yes | **** | <0.0001 |
| RJ vs. LPS RJEV 100 | -78.02 | Yes | *** | 0.0001 |
| RJ vs. LPS RJEV 1000 | -55.05 | Yes | * | 0.0252 |
| RJ vs. LPS RJEV 2500 | 18.91 | No | ns | >0.9999 |
| LPS RJEV 100 vs. LPS RJEV 1000 | 22.97 | No | ns | >0.9999 |
| LPS RJEV 100 vs. LPS RJEV 2500 | 96.93 | Yes | **** | <0.0001 |
| LPS RJEV 1000 vs. LPS RJEV 2500 | 73.96 | Yes | *** | 0.0004 |
| **10 to 14.5 hours** |  |  |  |  |
| **Dunn's multiple comparisons test** | **Mean rank diff.** | **Significant?** | **Summary** | **Adjusted P Value** |
| ctrl vs. LPS | -125 | Yes | **** | <0.0001 |
| ctrl vs. RJ | 32.97 | No | ns | 0.8608 |
| ctrl vs. LPS RJEV 100 | -50.5 | No | ns | 0.0541 |
| ctrl vs. LPS RJEV 1000 | -16.39 | No | ns | >0.9999 |
| ctrl vs. LPS RJEV 2500 | 34.75 | No | ns | 0.6777 |
| LPS vs. RJ | 157.9 | Yes | **** | <0.0001 |
| LPS vs. LPS RJEV 100 | 74.47 | Yes | *** | 0.0003 |
| LPS vs. LPS RJEV 1000 | 108.6 | Yes | **** | <0.0001 |
| LPS vs. LPS RJEV 2500 | 159.7 | Yes | **** | <0.0001 |
| RJ vs. LPS RJEV 100 | -83.47 | Yes | **** | <0.0001 |
| RJ vs. LPS RJEV 1000 | -49.36 | No | ns | 0.0666 |
| RJ vs. LPS RJEV 2500 | 1.78 | No | ns | >0.9999 |
| LPS RJEV 100 vs. LPS RJEV 1000 | 34.11 | No | ns | 0.7394 |
| LPS RJEV 100 vs. LPS RJEV 2500 | 85.25 | Yes | **** | <0.0001 |
| LPS RJEV 1000 vs. LPS RJEV 2500 | 51.14 | Yes | * | 0.048 |
| **10 to 14.5 hours** |  |  |  |  |
| **Dunn's multiple comparisons test** | **Mean rank diff.** | **Significant?** | **Summary** | **Adjusted P Value** |
| ctrl vs. LPS | -89.81 | Yes | **** | <0.0001 |
| ctrl vs. RJ | 36.64 | No | ns | 0.5204 |
| ctrl vs. LPS RJEV 100 | -43.47 | No | ns | 0.1834 |
| ctrl vs. LPS RJEV 1000 | -6.26 | No | ns | >0.9999 |
| ctrl vs. LPS RJEV 2500 | 59.58 | Yes | ** | 0.0089 |
| LPS vs. RJ | 126.5 | Yes | **** | <0.0001 |
| LPS vs. LPS RJEV 100 | 46.34 | No | ns | 0.1134 |
| LPS vs. LPS RJEV 1000 | 83.55 | Yes | **** | <0.0001 |
| LPS vs. LPS RJEV 2500 | 149.4 | Yes | **** | <0.0001 |
| RJ vs. LPS RJEV 100 | -80.11 | Yes | **** | <0.0001 |
| RJ vs. LPS RJEV 1000 | -42.9 | No | ns | 0.2011 |
| RJ vs. LPS RJEV 2500 | 22.94 | No | ns | >0.9999 |
| LPS RJEV 100 vs. LPS RJEV 1000 | 37.21 | No | ns | 0.4796 |
| LPS RJEV 100 vs. LPS RJEV 2500 | 103.1 | Yes | **** | <0.0001 |
| LPS RJEV 1000 vs. LPS RJEV 2500 | 65.84 | Yes | ** | 0.0022 |
| **15 to 19.5 hours** |  |  |  |  |
| **Dunn's multiple comparisons test** | **Mean rank diff.** | **Significant?** | **Summary** | **Adjusted P Value** |
| ctrl vs. LPS | -61.28 | Yes | ** | 0.003 |
| ctrl vs. RJ | 31.48 | No | ns | 0.8379 |
| ctrl vs. LPS RJEV 100 | -19.66 | No | ns | >0.9999 |
| ctrl vs. LPS RJEV 1000 | 14.64 | No | ns | >0.9999 |
| ctrl vs. LPS RJEV 2500 | 60.61 | Yes | ** | 0.0035 |
| LPS vs. RJ | 92.76 | Yes | **** | <0.0001 |
| LPS vs. LPS RJEV 100 | 41.62 | No | ns | 0.1719 |
| LPS vs. LPS RJEV 1000 | 75.92 | Yes | **** | <0.0001 |
| LPS vs. LPS RJEV 2500 | 121.9 | Yes | **** | <0.0001 |
| RJ vs. LPS RJEV 100 | -51.13 | Yes | * | 0.0284 |
| RJ vs. LPS RJEV 1000 | -16.83 | No | ns | >0.9999 |
| RJ vs. LPS RJEV 2500 | 29.13 | No | ns | >0.9999 |
| LPS RJEV 100 vs. LPS RJEV 1000 | 34.3 | No | ns | 0.558 |
| LPS RJEV 100 vs. LPS RJEV 2500 | 80.27 | Yes | **** | <0.0001 |
| LPS RJEV 1000 vs. LPS RJEV 2500 | 45.97 | No | ns | 0.0785 |
| **Growth rate** |  |  |  |  |
| **Tukey's multiple comparisons test** | **Mean Diff.** | **95.00% CI of diff.** | **Summary** | **Adjusted P Value** |
| ctrl vs. LPS | -0.455 | -1.478 to 0.5679 | ns | 0.7407 |
| ctrl vs. RJ | 0.04167 | -0.9813 to 1.065 | ns | >0.9999 |
| ctrl vs. LPS RJEV 100 | 0.2215 | -0.8014 to 1.244 | ns | 0.9837 |
| ctrl vs. LPS RJEV 1000 | -0.02995 | -1.053 to 0.9930 | ns | >0.9999 |
| ctrl vs. LPS RJEV 2500 | -0.2129 | -1.236 to 0.8100 | ns | 0.9863 |
| LPS vs. RJ | 0.4967 | -0.5263 to 1.520 | ns | 0.6667 |
| LPS vs. LPS RJEV 100 | 0.6766 | -0.3464 to 1.700 | ns | 0.3479 |
| LPS vs. LPS RJEV 1000 | 0.4251 | -0.5979 to 1.448 | ns | 0.7902 |
| LPS vs. LPS RJEV 2500 | 0.2421 | -0.7808 to 1.265 | ns | 0.9759 |
| RJ vs. LPS RJEV 100 | 0.1799 | -0.8431 to 1.203 | ns | 0.9936 |
| RJ vs. LPS RJEV 1000 | -0.07162 | -1.095 to 0.9513 | ns | >0.9999 |
| RJ vs. LPS RJEV 2500 | -0.2546 | -1.278 to 0.7684 | ns | 0.9701 |
| LPS RJEV 100 vs. LPS RJEV 1000 | -0.2515 | -1.274 to 0.7715 | ns | 0.9716 |
| LPS RJEV 100 vs. LPS RJEV 2500 | -0.4345 | -1.457 to 0.5885 | ns | 0.7751 |
| LPS RJEV 1000 vs. LPS RJEV 2500 | -0.183 | -1.206 to 0.8400 | ns | 0.9931 |
